# A mosaic RBD-nanoparticle pan-sarbecovirus vaccine elicits nanobodies that target highly conserved epitopes

**DOI:** 10.64898/2026.10.13.758647

**Authors:** Kathryn E. Malecek, Jennifer R. Keeffe, Chengcheng Fan, Jaasiel Alvarez, Alyssa Player, Han Gao, Priyanthi P.N.P. Gnanapragasam, Semi Rho, Lisa Schimanski, Luisa N. Segovia, Tiong Kit Tan, Pamela J. Bjorkman

## Abstract

SARS-CoV-2, the virus that causes COVID-19, continually mutates, producing new variants of concern (VOCs). Substitutions in the SARS-CoV-2 spike protein receptor-binding domain (RBD) have resulted in reduced efficacies of vaccines and therapeutic monoclonal antibodies. Indeed, since the emergence of Omicron and earlier VOCs, most previously approved anti-RBD therapeutic monoclonal antibodies are no longer authorized as therapeutics. To address VOC substitutions effects on therapeutic antibody efficacy, we sought to identify single-domain nanobodies (VHHs) targeting RBD regions (class 4, class 1/4, portions of class 3 and class 5 RBD epitopes) that are more conserved across VOCs, reasoning that the smaller size of camelid nanobodies compared with conventional two-domain antibody combining sites would facilitate access to occluded RBD epitopes. We used a Beacon optofluidic platform to isolate and characterize single memory B cells from a llama immunized with a mosaic RBD-nanoparticle immunogen that elicits broad antibody responses across sarbecoviruses. Using multiplex binding assays, we identified nanobodies that bind to spike RBDs from diverse sarbecovirus strains. Competition ELISAs and crystal structures of VHH-RBD complexes showed that isolated nanobodies recognize broadly conserved epitopes, including those on the RBD that are inaccessible to analogous conventional two-domain antibodies. Promising VHH-Fcs neutralized SARS-CoV-2 variants of concern and zoonotic sarbecoviruses, demonstrating potential therapeutic utility for current and future outbreaks.

## Introduction

Spillover of zoonotic SARS-like betacoronaviruses (sarbecoviruses) resulted in two human health emergencies in the past 25 years: the SARS-CoV (hereafter SARS-1) epidemic in the early 2000s and the more recent COVID-19 pandemic caused by SARS-CoV-2 (hereafter SARS-2). The existence of large coronavirus reservoirs in bats raises the possibility of cross-species transmission that could result in future epidemics or pandemics.^1–3^ In addition, SARS-2 variants emerged throughout the COVID-19 pandemic, with the Alpha, Beta, Delta, Gamma, and Omicron lineages designated as variants of concern (VOCs) due to apparent increased transmissibility and/or resistance to neutralizing antibodies (Abs) elicited by infection or vaccination.^4–7^ In the case of Omicron VOCs, the many substitutions in the SARS-2 spike protein receptor-binding domain (RBD), the major target of neutralizing Abs^8–14^ and detectable cross-variant neutralization,^15^ resulted in reduced efficacies of vaccines and therapeutic monoclonal Abs (mAbs).^7,16^ Comparison of the variability of RBDs across sarbecoviruses and within SARS-2 VOCs suggest that vaccines and mAbs targeting the more conserved neutralizing Ab epitopes (class 4, class 1/4, and portions of class 3 and class 5 epitopes^17–20^) (Figure 1A) could protect against present and future SARS-2 VOCs. By contrast, Abs targeting less conserved class 1 and class 2 RBD epitopes that directly overlap with the binding footprint for human ACE2, the SARS-2 host receptor, recognize a portion of the RBD that exhibits sequence variability between sarbecoviruses.^17^

**Figure 1.**
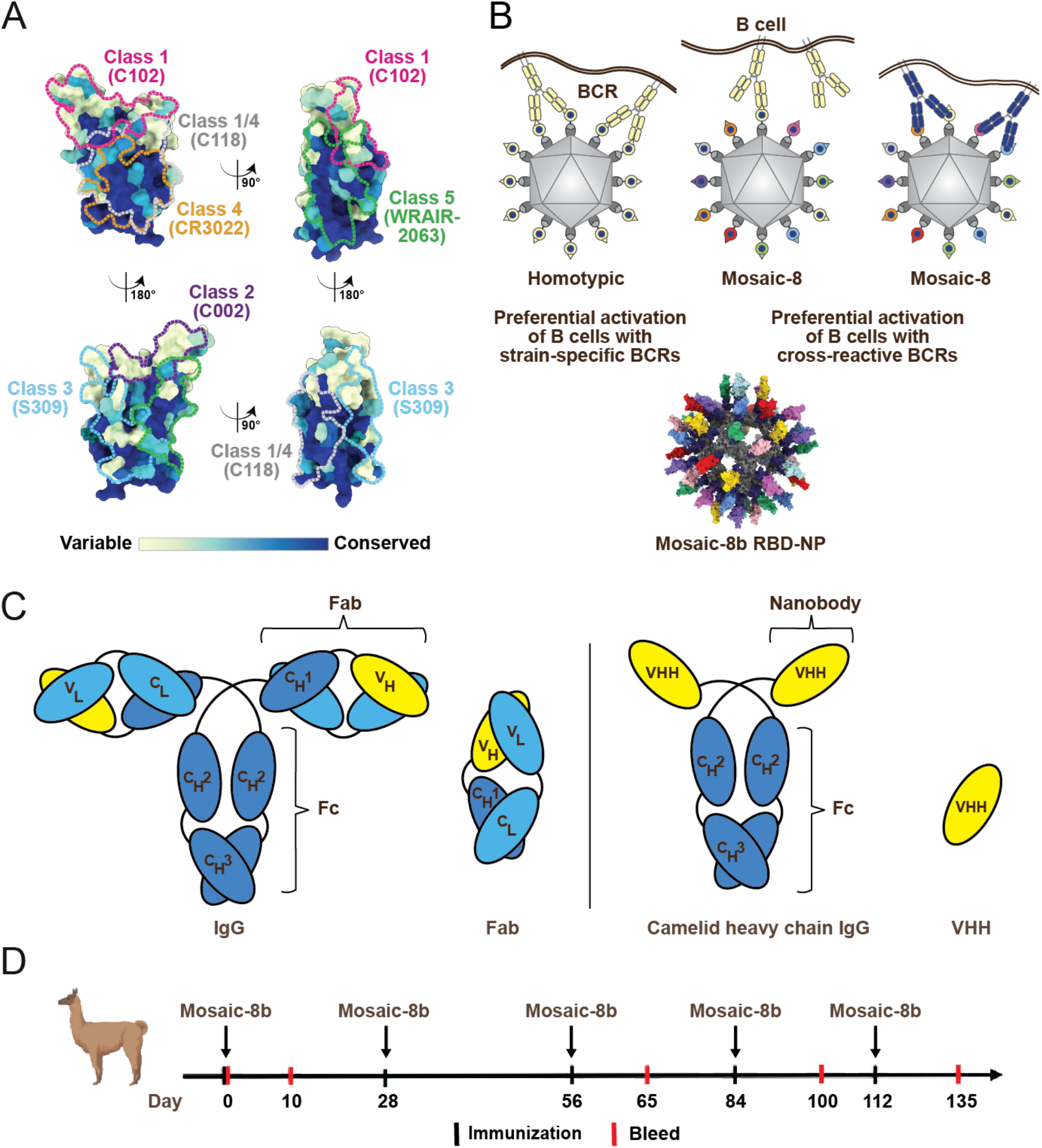
Strategy to induce Abs against conserved RBD epitopes. (A) Surface representation of SARS-2 RBD (PDB 7BZ5) showing sequence conservation of 16 sarbecovirus RBDs (calculated using ConSurf^74^). Anti-RBD Ab epitopes recognizing class 1, 2, 3, 4, 5 and 1/4 RBD epitopes^17–20^ are outlined in dots in different colors using information from structures of Abs bound to SARS-2 spike or RBD (C102: PDB 7K8M; C002: PDB 7K8T, S309: PDB 7JX3; CR3022: PDB 7LOP; C118: PDB 7RKV; WRAIR-2063: PDB 8EOO). (B) Top: Avidity hypothesis. Left: Strain-specific BCRs (pale yellow) utilizing avidity to recognize a strain-specific epitope (pale yellow triangle) on identical antigens attached to a homotypic NP. Middle: Strain-specific BCRs binding with only one Fab (not using avidity) to a strain-specific epitope (triangle) on an antigen attached to a mosaic NP. Right: Cross-reactive BCRs using avidity to recognize a common epitope (blue circle) presented on different antigens attached to a mosaic NP, but not to strain-specific epitopes (triangles). Not all 60 attached RBDs on NPs are shown for clarity. Bottom: Model of mosaic-8b RBD-NP made using coordinates of an RBD (PDB 7SC1) (different colors), mi3 (PDB 7B3Y) (gray), and SpyCatcher (PDB 4MLI) (blue). (C) Comparison of conventional and camelid IgGs. Left: Conventional IgG and Fab. Antigen recognition is conferred by a heterodimer comprising a paired variable heavy domain and variable light domain (VH-VL). Right: Heavy chain-only camelid IgG with its antigen-binding VHH domain. Antigen recognition is conferred by a single monomeric domain, which has greater access to occluded regions of a coronavirus spike trimer. (D) Immunization regimen. A llama was immunized with 55 µg (based on RBD content) of mosaic-8b RBD-NPs at days 0, 28, 56, 84, and 112, and blood samples were collected at days 10, 65, 100, and 135.

We previously hypothesized that B cell receptor (BCR) crosslinking between adjacent conserved epitopes on RBDs presented on a nanoparticle (NP) would preferentially activate B cells that make Abs against more conserved regions of RBDs compared with B cells producing Abs against immunodominant variable regions^21^ (Figure 1B). We subsequently showed that a 60-mer protein NP presenting 8 different RBDs (mosaic-8b RBD-NP; Figure 1B), elicited potent and broadly cross-reactive mAbs in mice and rabbits that are predicted to retain potency upon emergence of new SARS-2 VOCs or another sarbecovirus spillover event.^22,23^ By contrast, both infection and available COVID vaccinations tend to elicit strain-specific class 1 and class 2 anti-RBD Abs that lose potency when a new SARS-2 VOC emerges.^24–26^ Indeed, since the emergence of Omicron and earlier VOCs, class 1, 2, and 3 anti-RBD therapeutic mAbs that were previously approved for COVID-19 treatment (e.g., bamlanivimab plus etesevimab, casirivimab plus imdevimab, and sotrovimab) are no longer authorized as therapeutics due to reduction in their activity against VOCs.^27^ At present, only one anti-SARS-2 mAb, Pemgarda, is authorized by the FDA for use in humans^28^ but this mAb recognizes a relatively variable class 1 RBD epitope.^23,29^ There are currently no approved mAb therapeutics targeting more conserved RBD epitopes. mAbs against more conserved epitopes (class 4, class 1/4, and class 5) are rare after infection or immunization, possibly because these epitopes are somewhat occluded within the coronavirus spike trimer.^18,22^ In addition, mAbs against conserved epitopes, while broad, often exhibit lower potencies.^21^ Thus, there remains a need for therapeutic mAbs that recognize more conserved RBD epitopes and retain activity despite continued viral evolution.

Broadly cross-reactive Abs that are smaller than conventional IgGs should be better suited to access conserved occluded epitopes at the RBD base. Whereas conventional mammalian IgGs are composed of two heavy chains and two light chains arranged into two variable heavy-variable light (VH-VL) domain-containing Fabs and a dimeric C-terminal Fc region, llamas and related camelids produce single-domain heavy-chain only IgGs (HC-IgGs), in which a single-domain nanobody (VHH) is linked to each chain of the dimeric Fc (Figure 1C). Owing to the absence of light chains and the constant heavy domain 1 (CH1) in the Fab, VHHs are smaller than a conventional VH-VL Fab and exhibit high stability,^30^ making both natural and engineered VHHs valuable tools in therapeutics and biotechnology.^31^ We hypothesized that the smaller size of VHHs might facilitate access to conserved RBD epitopes that are sterically occluded on the spike trimer and therefore less accessible to conventional IgGs. We therefore investigated whether HC-IgGs against the more conserved RBD epitopes could be elicited by immunizing a llama with mosaic-8b RBD-NPs (Figure 1D) that preferentially elicit Abs against more conserved RBD epitopes.^21–23,32–34^ We used single-cell optofluidics and multiplexed antigen specificity assays to characterize antigen recognition and evaluate the cross-reactivity of IgGs elicited by mosaic-8b. By assessing neutralization potencies against sarbecoviruses and mapping elicited VHH epitopes using competition ELISAs and 3D structure determinations, we demonstrated that neutralizing VHHs that target conserved epitopes can be elicited in a llama. Notably, the majority of the HC-IgGs that we characterized recognized conserved RBD epitopes, consistent with the hypothesis that the smaller size of HC-IgGs enables access to epitopes that are less accessible to conventional IgGs.

Together, these findings establish a rapid and high-throughput optofluidics-based strategy for the discovery of antigen-specific camelid HC-IgGs. The HC-IgGs identified here after immunization of a llama with a pan-sarbecovirus vaccine candidate represent potentially valuable therapeutic reagents with efficacy against emerging SARS-2 VOCs and future zoonotic sarbecovirus spillovers.

## Materials and Methods

### Mosaic-8b RBD-NP llama immunizations

Mosaic-8b RBD-NPs were produced using SpyTagged RBDs expressed in *Pichia pastoris* and SpyCatcher003-mi3 NPs^35^ expressed in *B. subtilis*.^33^ Concentrations of conjugated RBD-NPs (determined using a Bio-Rad Protein Assay) are reported based on RBD content. Mosaic-8b RBD-NP immunizations were performed at the University of Reading animal facility using IACUC-approved protocols. A llama (*Lama glama*) was immunized intramuscularly with 55 µg (based on RBD content) of mosaic-8b RBD-NPs in Tris-buffered saline (TBS) mixed 1:1 (v/v) with GERBU F adjuvant (Biotechnik GmbH, catalog number: 3030) at days 0, 28, 56, 84, and 112, following previous protocols for repeated immunizations in llamas and under appropriate approval by the University of Reading IACUC.^36^ PBMCs and serum were isolated from blood samples collected at days 10, 65, 100, and 135 (Figure 1D). To isolate PBMCs, an equal amount of fresh whole blood was added to sterile PBS and mixed. 15 mL of Lymphoprep (StemCell Technologies) was added into a SepMate50 tube (StemCell Technologies), and 30 mL of the diluted whole blood was added. The tube was spun at 1,200 x g for 10 min at room temperature (RT). Isolated PBMCs and serum were then transferred into a 50 mL Falcon tube, to which sterile PBS was added to 50 mL and then centrifuged at 300 x g for 8 min to wash the lymphocytes. The washing step was repeated twice. On the last wash, the tube was spun at 120 x g for 10 min with a brake on. The supernatant was then carefully discarded, leaving the PBMC pellet secure at the bottom of the tube. PBMCs were resuspended in freezing media (Fetal Bovine Serum with 10% DMSO) and stored in LN2 until further downstream processing. To isolate serum, whole blood collected into red-capped Vacutainer blood collection tubes (BD) and incubated at RT for ≥1 hour. Tubes were then centrifuged for 10 min at 1,400 x g, and serum was then aliquoted into clean tubes for storage.

### Recombinant protein expression and purification

Recombinant proteins, including VHHs, VHH-Fcs (VHH fused to a human IgG1 Fc), RBDs, and spike trimers, were expressed via transient transfection of Expi293 or Expi293 PRO cells (Gibco) using ExpiFectamine or Expi293 PRO Transfection Reagent, respectively. Biotinylated RBDs were prepared by co-expression of an Avi-tagged RBD with BirA enzyme for in vivo biotinylation (kind gift of Michael Anaya, Caltech) using a 1:1 RBD to BirA plasmid ratio for transfections and then purified from transfection supernatants by nickel affinity chromatography (HisTrap HP, Cytiva). VHH-Fc constructs and IgG controls were purified from transfection supernatants using MabSelect SuRe columns (Cytiva). RBDs and spike trimers for ELISAs were purified from transfection supernatants by nickel affinity chromatography (HisTrap HP, Cytiva), followed by size exclusion chromatography (Superdex 200 Increase 10/300 GL, Cytiva). VHHs and RBDs for crystallographic studies were co-expressed by transient transfection of VHH- and 6xHis-tagged RBD-encoding expression plasmids and complexes were purified from transfected cell supernatants using nickel affinity chromatography and size exclusion chromatography (Superdex 200 Increase 10/300 GL, Cytiva). Fractions corresponding to VHH-RBD complexes were pooled, concentrated to ∼20 mg/mL, and stored at 4°C.

XVR011 VHH-Fc (rimteravimab; an affinity matured version of VHH-72^37^) was obtained from MedChemExpress (catalog number HY-P99947). The sequence of XVR011 was inferred from descriptions of changes made to the published sequence of VHH-72.^38^

### Bruker Cellular Analysis Beacon Assays

Memory B cells expressing B cell receptors (BCRs) that bind RBD were enriched by incubating 7 x 10^6^ llama PBMCs from Day 65 and Day 96 with 2 µg each of five biotinylated RBDs (matched with respect to mosaic-8b: SHC014, Rs4081, SARS-2 Beta; and mismatched: Khosta-2 and SARS-CoV) followed by pulldown using magnetic-activated cell sorting (MACS) with streptavidin-coated microbeads (Miltenyi 130-105-637). A PE-conjugated anti-alpaca IgG2/3 Ab (Bruker Cellular Analysis KT52-00007) followed by pulldown with anti-PE microbeads (Miltenyi 130-048-801) were then used to enrich cells expressing IgG2/3 subclass BCRs. Recovered memory B cells were incubated in activation culture media (Bruker Cellular Analysis KT52-00007) at 2.5 x 10^4^ cells/mL in a V-bottom plate. Over 5 days, the isolated memory B cells were expanded and activated to secrete IgG for assay purposes according to the manufacturer’s protocol (Bruker Cellular Analysis, catalog number KT520010) to promote differentiation into plasmablasts.

Activated memory B cells were loaded onto the Beacon instrument and isolated into individual nanopen culture compartments as described.^23^ Assays to detect secreted IgG were performed using 5% (w/v) slurries of Bruker Anti-Camelid IgG (H+L) Assay Beads (Bruker Cellular Analysis RG52-00026) or Spherotech protein G-coated particles (PGP-60-5) to capture secreted Abs on beads. The secreted captured Ab was detected with either a labeled anti-IgG secondary Ab for total IgG (Alexa Fluor 647-conjugated goat anti-alpaca IgG (H+L) Jackson Immuno Research 128-605-160, 1:1,000 dilution), or with anti-IgG2/3 subclass specific secondary Ab (Alexa Fluor 488 AffiniPure goat anti-alpaca IgG subclasses 2+3 Specific, Jackson Immuno Research 128-545-229, 1:1,000 dilution). Fluorescently labeled Fabs were used to detect the accumulation of secreted Ab on the Protein G beads (Alexa Fluor 488-conjugated AffiniPure Fab fragment, Jackson Immuno Research 109-546-006, 1:500 dilution, or goat anti-alpaca IgG, VHH domain, Jackson Immuno Research 128-547-230, 1:500 dilution). Biotinylated RBDs were labeled with streptavidin-fluorophore conjugates, including Brilliant Violet 421-Streptavidin, Biolegend 405225SA-AF488, Thermo Scientific S32354, SA-AF555, Thermo Scientific S32355, and SA-AF680, Thermo-Scientific S32358. RBD-fluorophore complexes were included in multiplexed assays as soluble antigens (0.5 µg/mL; ∼20 nM). SpyCatcher003-mi3 was chemically biotinylated and immobilized on streptavidin-coated particles (Spherotech SVP-60-5) without fluorescent labeling and used as a 5% (w/v) slurry with Abs specific for the IgG2/3 subclass and IgG total to screen for NP scaffold-reactive Abs.

Nanopens containing single activated memory B cells were scored for IgG2/3 secretion and RBD reactivity. Cells were selected for export based on their RBD binding scores and IgG2/3 secretion scores. Single cells were exported into individual wells of a 96 well plate containing lysis buffer, and total RNA was isolated for single cell cDNA library preparation via reverse transcription (Bruker Cellular Analysis 750-02030). PCR products were generated using a forward primer specific to the 5’ library handle and a reverse primer that binds within a sequence encoding the CH2 domain of llama IgG subclass genes (Table S1). PCR products were then sequenced, and the hinge regions were analyzed to determine the IgG subclass. A cell was determined to encode a HC-IgG if the hinge region sequence was consistent with the annotated IgG2 or IgG3 subclass^39,40^ and a CH1 region-encoding sequence was not observed. VHH-encoding sequences were cloned from HC-IgG cells into a human IgG1 Fc (IgG1 hinge and CH2-CH3 domains) expression vector using the primers listed in Table S1 to generate VHH-Fcs.

VHH sequences were annotated and analyzed using PipeBio (Benchling) software using the sequences of llama Ab germline gene segments using IMGT (Table S2).^41^ VHH protein sequence alignments were conducted using MUSCLE^42^ to generate phylogenetic trees and pairwise identity heatmaps.

### Binding and Competition ELISAs

Serum ELISAs were performed by coating purified His-tagged RBD (2 μg/mL in phosphate-buffered saline; PBS) onto 96-well ELISA plates (Thermo Fisher Scientific) and incubating at room temperature for 1 hour. Coated plates were then washed 3 times with PBS and blocked with 5% skim milk in PBS over night at 4°C. Heat-inactivated (56°C for 30 min) llama serum (1:10 initial dilution) was serially diluted by 2-fold in PBS/0.1% bovine serum albumin (BSA) and then added to the coated plates for 1 hour at room temperature, followed by washing with PBS, incubating for 1 hour with a 1:5,000 dilution of horse radish peroxidase (HRP)-conjugated rabbit anti-camelid secondary (GenScript, A02016-200), and then washing with PBS. Plates were then developed with KPL SureBlue TM TMB substrate (Sera Care, 5120-0075) for 5 minutes, and the reaction was stopped with 1M H2SO4. Absorbance was measured using a ClarioSTAR plate reader (BMG Labtech). All samples were assayed in duplicate, and data were analyzed and plotted using GraphPad Prism v10.6.1.

VHH-Fc ELISAs were performed using a Tecan Evo liquid handling robot as previously described.^33^ RBDs or spike trimers were adsorbed overnight to 384-well Maxisorp plates (Nunc) at 4 °C, then blocked with 3% BSA (w/v) in TBS-T (TBS containing 0.1% Tween-20 [v/v]) for 1 hour at room temperature. Purified VHH-Fcs were serially diluted in blocking solution, added to plates, and incubated for 3 hours at room temperature. Plates were washed with TBS-T and then incubated for 1 hour with a 1:200,000 dilution of HRP-conjugated anti-IgG secondary (goat anti-human IgG; SouthernBiotech 2015-05). Wells were washed with TBS-T and SuperSignal™ ELISA Femto Substrate (ThermoFisher 37074) was added as per manufacturer’s instructions prior to measuring luminescence at 425 nm. Data were collected in duplicate and midpoint titers (ED50 values) were obtained using GraphPad Prism v11.0.2 using a four-parameter dose-response curve fit. Data presented are representative of at least two independent experiments.

Competition ELISAs were performed using the Tecan Evo liquid handling robot as described.^22,23^ To identify the probable epitopes of VHH-Fc proteins, a Fab corresponding to a characterized IgG of known epitope was adsorbed to a 384-well Maxisorb ELISA plate, followed by blocking with 3% BSA (w/v) in 1x TBS-T. SARS-2 WA1 RBD was captured, and then 1 µg/mL VHH-Fc or control IgG was added to each well. Bound VHH-Fc or IgG was detected using HRP-conjugated goat anti-human Fc (Southern Biotech 2014-05) and SuperSignal ELISA Femto Substrate (ThermoFisher 37074). Average relative light units (RLUs) were plotted for measurements performed in quadruplicate using GraphPad Prism v11.0.2. Data presented are representative of at least two independent experiments.

### Pseudovirus Neutralization Assays

Lentiviral-based pseudoviruses were prepared in HEK293T cells as described^11,43^ using spike gene sequences encoding deletions of 21 residues (SARS-2 variants, WIV1, and SHC014) or 19 residues (SARS-1, Khosta-2-SARS-1 chimera, BtKY72-SARS-1 chimera) in the cytoplasmic tail. BtKY72 (including K493Y/T498W substitutions) and Khosta-2 pseudoviruses included chimeric spikes in which the RBD from SARS-1 (residues 323-501) was substituted with the RBD from BtKY72 K493Y/T498W (residues 327-503) or Khosta-2 (residues 324-500).^44^ Neutralization assays were done as described.^23^ Briefly, serially diluted VHH-Fc was incubated with pseudovirus for 1 hour at 37°C, then added to HEK293T-hACE2 target cells or for SHC014 assays, a high-hACE2 HEK-293T cell line expressing human ACE2 encoded with a consensus Kozak sequence (kindly provided by Kenneth Matreyek, Case Western Reserve University). The assay was incubated for 48 hours at 37°C. Cells were then lysed with Britelite Plus reagent (Revvity Health Sciences), and luminescence was measured. Each sample was tested in duplicate, and percent neutralization was plotted as a function of VHH-Fc concentration. Average half-maximal inhibitory dilutions (ID50) were derived in AntibodyDatabase^45^ using 4-parameter nonlinear regression. Data presented are representative of at least two independent experiments.

### X-ray crystallography

Crystallization trials were done at room temperature using the sitting drop vapor diffusion method with commercial screens by mixing 200 nL of VHH-RBD complex and 200 nL of crystallization solution using the MRC 2 well crystallization plate (Hampton Research) on a SPT LabTech Mosquito Instrument. Crystals for the VHH-RBD complexes were obtained under the following conditions: PEGRx screen (Hampton Research) containing 0.02 M nickel (II) chloride hexahydrate, 0.02 M magnesium chloride hexahydrate, 0.02 M cadmium chloride hydrate, 0.1 M sodium acetate trihydrate pH 4.5, and 24% (w/v) polyethylene glycol monomethyl ether 2,000 (P1-F6–RBD); PEG/Ion screen (Hampton Research) containing 2% (v/v) Tacsimate^TM^ pH 6.0, 0.1 M Bis-Tris pH 6.5, and 20% (w/v) polyethylene glycol 3,350 (P1-G11–RBD); JCSG Plus screen (Hampton Research) containing 0.1 M HEPES pH 7.5, 10% (w/v) polyethylene glycol 8,000, and 8% (v/v) ethylene glycol (P2-B5–RBD); JCSG Plus screen (Hampton Research) containing 0.2 M zinc acetate dihydrate, 0.1 M imidazole pH 8.0, and 20% (w/v) polyethylene glycol 3,000 (P2-G5–RBD). Crystals were cryoprotected in well solution with 25% glycerol before flash-cooling in liquid nitrogen.

X-ray diffraction datasets were collected at the Stanford Synchrotron Radiation Lightsource (SSRL) beamline 12-1 with an EIGER 2XE 16M PAD (Dectris) using the Blu-ice interface.^46^ Datasets were indexed and integrated with XDS^47^ and scaled with Aimless.^48^ VHH-RBD complex structures were solved by molecular replacement using coordinates of a SARS-2 RBD (PDB 7UZC) and Alphafold 3^49^ models of VHHs as inputs for *Phaser* in Phenix.^50^ VHHs were numbered consecutively for deposition and were labeled using the IMGT numbering scheme in Figure S4.^41^ Multiple refinement and model building cycles were carried out in *phenix.refine* in Phenix^50^ and *Coot.*^51^ Crystallographic refinement statistics for structures are reported in Table S3.

### Structure analyses

Structure figures were made using ChimeraX.^52^ VHH-RBD interfaces were assigned by identifying residues within 4 Å using PDBePISA.^53^ Hydrogen bonds were identified at VHH-RBD interfaces for acceptor/donor pairs using a distance criterion of ≤3.9 Å and an A-D-H angle >90°. van der Waals interactions were assigned using a maximum distance cutoff of 4.0 Å between non-hydrogen atoms.

### Modeling of VHH-spike and Fab-spike structures

Structures of SARS-2 spike in complex with P1-F6 VHH, P1-G11 VHH, P2-B5 VHH, and P2-G5 VHH were modeled by superimposing the coordinates of the RBD from a VHH-RBD complex structure onto the “up” RBDs of a SARS-2 spike (PDB: 7RKV) (Class 4/5 and class 4 VHHs could not be accommodated on RBDs in a “down” conformation; data not shown). To address whether a VH-VL–containing Fab recognizing the same RBD epitope on a spike trimer as a VHH, we superimposed the Cα coordinates of the VH domain of the human mAb C118 (PDB: 7RKV) onto the VHH Cα coordinates of a VHH-spike trimer model using UCSF ChimeraX^52^ and then asked whether steric clashes were predicted between the spike trimer and any of the four Fab domains (VH, VL, CH1, or CL). Steric clashes were inferred if atoms in a modeled Fab overlapped with atoms in the spike.

### QUANTIFICATION AND STATISTICAL ANALYSIS

Group size, replicates, and statistical analysis details can be found in the main text, figure legends, and method details.

## Results

### Characterization of llama antigen-reactive memory B cells using Beacon

Llamas produce VHH-containing IgG Abs that lack light chains due to a splicing defect mutation that disrupts inclusion of the exon encoding the CH1 domain^54^ and hydrophilic substitutions within IGHV genes at the counterpart of the interface with a VL domain.^55^ In camelids, these HC-IgGs are members of the IgG2 or IgG3 subclass. Llamas also produce IgG1 Abs that include light chains. To elicit HC-IgGs that recognize conserved epitopes on sarbecovirus RBDs, a llama (*Lama glama*) was immunized with mosaic-8b RBD-NP, a previously described pan-sarbecovirus RBD-NP vaccine candidate^21–23,33,34,56^ (Figure 1D). Mosaic-8b RBD-NPs were generated using the SpyCatcher-SpyTag system^57,58^ to randomly attach eight different SpyTagged RBDs (derived from SARS-2 Beta and seven other sarbecovirus RBDs from spikes in clade 1a, 1b, or 2 sarbecoviruses) to the 60 SpyCatchers on SpyCatcher-mi3 NPs.^59^ Mosaic-8b RBD-NPs were characterized and validated as previously described,^21–23,32,33,56,60^ and strain-specific RBD Abs^23^ were used to confirm the display of all 8 RBDs on the RBD-NPs (Figure 1D). Serum anti-RBD responses, as quantified by ELISA, were first observed starting at Day 65 (Figure S1). Peripheral blood mononuclear cells (PBMCs) were isolated from Day 65 and Day 100 blood samples (Figure 1D). Antigen-specific memory B cells were enriched by RBD pulldown followed by an IgG2/3 pulldown to enrich memory B cells that expressed HC-IgGs. Recovered cells were cultured to activate IgG secretion.

We used a Beacon optofluidics system (Figure 2A) to isolate and screen 13,656 single, activated memory B cells for secretion of IgGs from day 65 and day 100 PBMC samples that recognized diverse sarbecovirus RBDs from both matched strains (represented by an RBD on mosaic-8b) and mismatched strains (not present on mosaic-8b) (Figure 2B; Figure S2A). Assays for Ab subclass and antigen specificity were used to identify microfluidic chip culture compartments (nanopens) containing single memory B cells secreting an IgG that bound to a labeled antigen of interest. Serial multiplexed assays^23^ were performed using five fluorophores (Figure 2B; Figure S2A) to visualize fluorescently labeled Abs against IgG subclasses 2 and 3 or total IgG (heavy plus light chain; H+L) and binding of fluorescently labeled RBD antigens. Binding of each antigen or Ab was evaluated by the presence of fluorescent signal accumulation above an individual nanopen (Figure 3A). Signal was verified for assignment to a particular nanopen, quantified, and analyzed to determine binding breadth and subclass for each secreted IgG.

**Figure 2.**
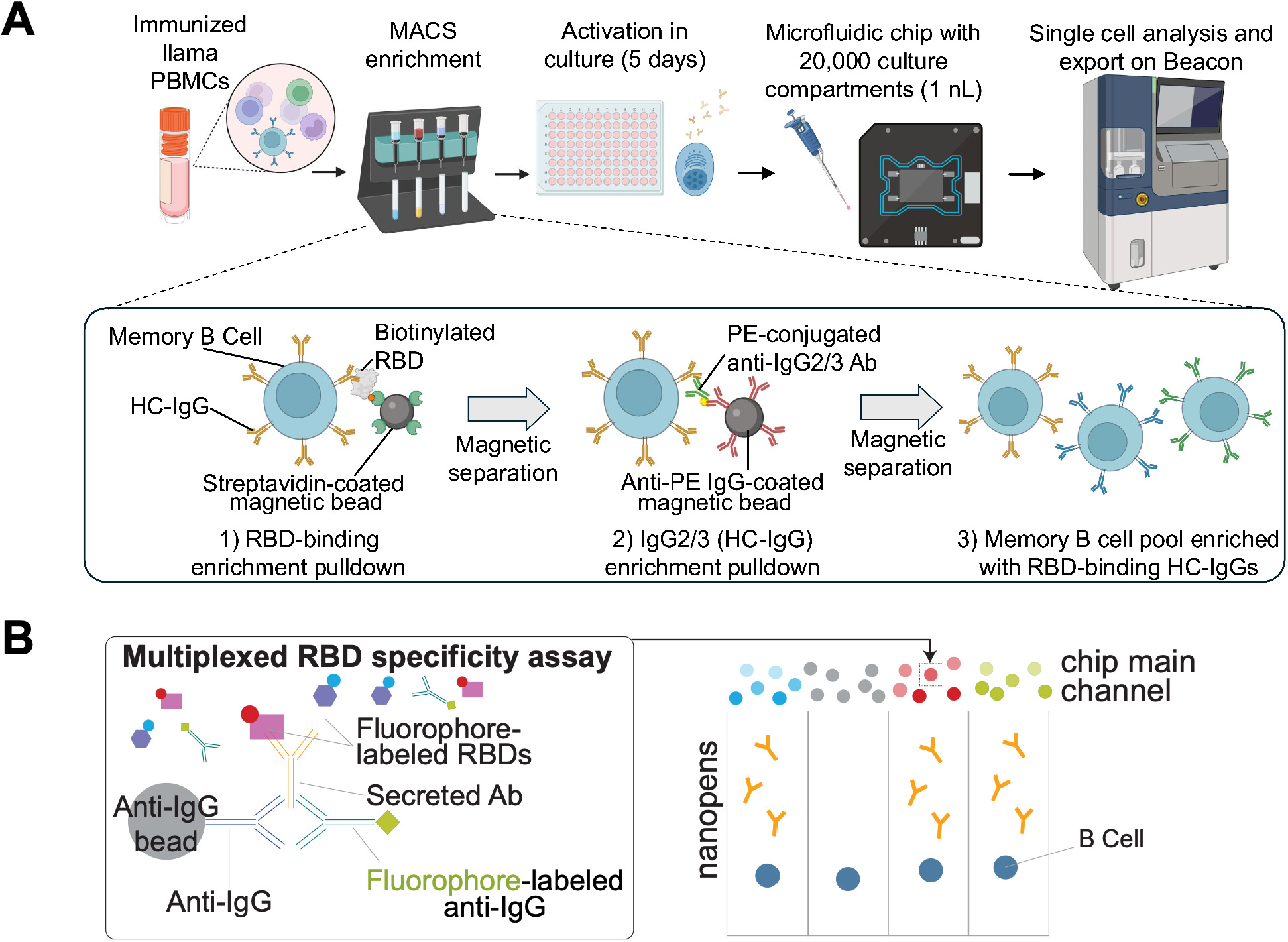
Enrichment of llama memory B cells for Beacon characterization. (A) Memory B cells displaying BCRs that recognize sarbecovirus RBDs were isolated via magnetic-activated cell sorting (MACS) enrichment. This pool of cells was further enriched for memory B cells displaying HC-IgGs. Recovered memory B cells were activated to secrete Abs in well plate culture and loaded onto a microfluidic chip with 20,000 nanopen culture compartments using the Beacon instrument. (B) Multiplexed antigen binding assays performed in a Beacon microfluidic chip. An assay bead mixture was prepared with camelid-specific IgG capture beads, fluorescently labeled soluble RBDs, and a fluorescently labeled anti-IgG secondary Ab. Images were acquired over time. Ab secretion and RBD binding were visualized as fluorescent signal accumulation above the nanopen containing an Ab-secreting cell.

**Figure 3.**
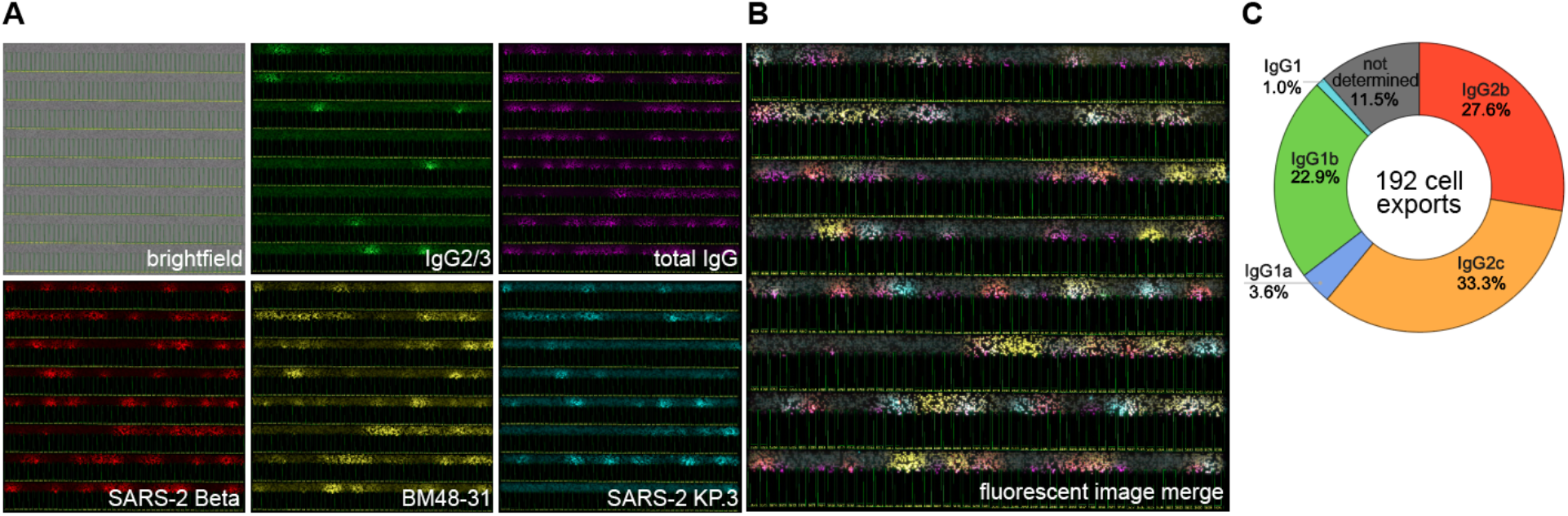
Multiplexed antigen specificity assays demonstrate broad RBD binding by Abs elicited in an immunized llama. Example of a multiplexed antigen binding assay using activated llama memory B cells, three soluble RBD antigens, and Abs for IgG2/3 subclasses or total IgG. Brightfield image shows assay beads in the main channel, and nanopen compartments, outlined in green and numbered in yellow. Cells are visible within nanopens. Images show one of the fifty fields of view on the chip, approximately 400 of the 20,000 nanopens. (A) Each channel shown individually. (B) Merged fluorescent image with five colors. For each assay, the signal above the nanopen was scored for each individual color and then analyzed together to profile the IgG subclass and RBD-binding properties of each isolated B cell. (C) Sequence recovery from exported cells by IgG subclass. The IgG hinge region was sequenced from each cell exported from the Beacon, showing enrichment for HC-IgG (IgG2) subclasses. IgG1 is reported when a sequence encoding CH1 is detected, but the exact IgG1 subtype cannot be determined.

As previously described for immunizations in mice and rabbits,^22,23^ we found that mosaic-8b immunization of a llama resulted in memory B cells that recognize RBDs from both matched and mismatched sarbecoviruses. 37.4% of IgG-positive nanopens were positive for ≥4 of 6 mismatched antigens evaluated on the Beacon, demonstrating the ability of mosaic-8b to elicit Ab responses to mismatched RBDs in a llama. IgG2-positive nanopens exhibited less binding breadth than IgG total positive/IgG2 negative nanopens (15.2% of IgG2+ nanopens bound ≥4 RBDs, 22.2% of IgG1/3+ nanopens bound ≥4 RBDs).

Cells were selected for export based on their binding breadth to RBDs and subclass determination for IgG2/3.^23^ We prioritized the export of cells that secreted Abs that showed recognition of (i) all six mismatched RBDs, (ii) at least one mismatched RBD, (iii) at least one (matched) RBD from clades 1a, 1b, and 2, and in addition, those that (iv) showed secretion of IgG2/3 HC-IgG. We isolated total RNA from the lysate of each exported cell, generated single cell cDNA libraries, and then amplified and sequenced the variable heavy (VH) and hinge regions of each IgG gene to determine the VH sequence and confirm the IgG subclass (Figure 3B; Figure S2B). VHH sequences were subcloned into an expression vector encoding the human IgG1 hinge and Fc to generate VHH-Fcs. We observed hinge and CH1 sequences consistent with IgG1a, IgG1b, IgG2b and IgG2c subclasses but no sequences corresponding to the IgG3 subclass, consistent with published results showing there are fewer IgG subclasses in llamas than in other camelids such as alpacas.^39^

Following confirmation of IgG subclasses by sequencing (Figure 3B), we revisited the Beacon assay data on subclass determination (Figure S2B). We found that Beacon assays correctly identified 90.6% IgG1 subclass sequences (signal with IgG total and not with IgG2/3), but only 52.1% of the HC-IgG sequences were correctly identified using IgG2/3 signal on the Beacon. The average bloom intensities for IgG2 were lower (below scoring threshold) for nanopens that were later determined to express HC-IgGs (false negatives on Beacon). This suggests lower secretion of IgG2 Abs and/or low sensitivity of the anti-IgG2/3 antibody used for detection in the multiplex assay format may cause false negatives for HC-IgG identification. This suggests that greater sensitivity, at the expense of some specificity, can be gained from filtering on the ratio of IgG2 and total IgG signals when analyzing Beacon data.

We also found five instances in which nanopens were identified as IgG2-positive but were determined by sequencing to contain IgG1 mAbs. In each case, adjacent nanopens to these false positive nanopens had modest IgG2 bloom scores. This suggests that local diffusion of a bloom signal can lead to misassignment of signals to nanopens. Collectively, our data show that sequence confirmation of IgG subclass is essential before proceeding to cloning for expression and validation of VHHs.

### Characterization of VHH-Fcs

RBD recognition by the recovered VHH-Fcs elicited by mosaic-8b RBD-NPs was assessed via ELISA against a larger panel of 20 RBDs that included 7 SARS-2 RBDs (WA1, Beta, BA.4/5, XBB.1.5, JN.1, LB.1, KP.3) and 13 animal sarbecovirus RBDs (RaTG13, Pang17, RshSTT200, SARS-1, WIV1, SHC014, Rs4081, RmYN02, Rf1, Yun11, BtKY72, Khosta-2, BM48-31), of which 8 RBDs were matched and 12 RBDs were mismatched with respect to the mosaic-8b immunogen. Nine VHH-Fcs recognized all antigens tested, and the remaining VHH-Fcs bound to RBDs across multiple clades (Figure 4A). The ELISA binding profiles, assessed using single RBD antigens, were generally consistent with what was observed in the single cell multiplexed assays on the Beacon (Figure S5A). In addition, we found that all VHH-Fcs recognized the SARS-2 RBD in the context of an intact spike trimer, suggesting they can access their epitope in a natively-folded conformation (Figure S5B).

**Figure 4.**
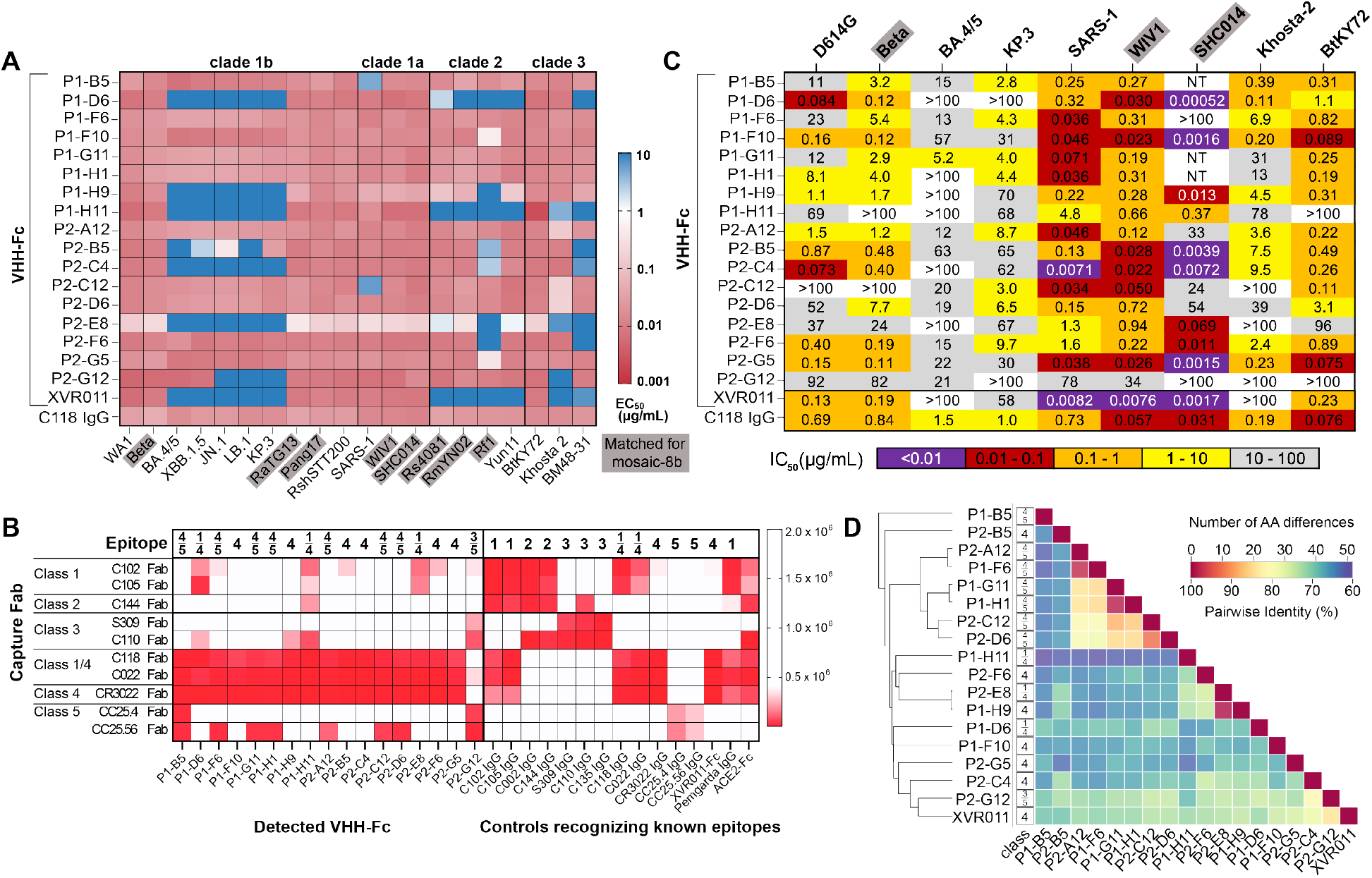
VHH-Fcs are broadly cross-reactive and recognize conserved RBD epitopes. (A) ELISA-derived EC50 values for recovered VHH-Fcs against a panel of sarbecovirus RBDs. RBDs that were included on mosaic-8b RBD-NPs (matched) are outlined in a gray box. Other listed RBDs are mismatched with respect to mosaic-8b. (B) Competition ELISA mapping of VHH epitopes. An immobilized Fab from a structurally characterized mAb (listed as Capture Fab; PDB codes for the Fab-RBD or Fab-spike structures are 7K8M, CXCM, 7K90, 6WPT, 7K8V, 7RKS, 7RKU, 6W41, 8SDF, and 8SDH) was used to capture SARS-2 RBD, a VHH-Fc, conventional IgG mAb, or ACE2-Fc (listed at the bottom) was added, and Fc was detected. White: high luminescence signal indicating IgG binding does not compete with the Fab. Red: low signal indicating reduced binding consistent with competition. The top line shows the RBD epitope class identified for each VHH-Fc or control protein based on the competition ELISA results. (C) Neutralization potencies of VHH-Fcs in a pseudovirus neutralization assay. Pseudoviruses containing RBDs included on mosaic-8b RBD-NPs (matched) are outlined in a gray box. (D) Phylogenetic tree relating VHHs (vertical, on left) shown with pairwise amino acid sequence identities.

We further characterized RBD epitopes recognized by elicited VHHs using a previously described competition ELISA^23^ (Figure 4B). Fabs generated from characterized human mAbs that recognize class 1, 2, 3, 4, 1/4, or 5 RBD epitopes were adsorbed on an ELISA plate and then incubated with SARS-2 WA1 RBD. VHH-Fcs derived from llama memory B cells or control IgGs of known epitopes were then added and evaluated for binding. A VHH-Fc or IgG will only bind to WA1 RBD if it is not sterically occluded by the bound Fab. We determined the probable epitopes of 19 VHH-Fcs isolated from llamas and two control VHH-Fcs from literature on SARS-2 WA1 RBD (Figure 4B). As expected for the degree of binding breadth observed on the Beacon and by ELISA, we identified 16 VHH-Fcs that competed with a class 4 anti-RBD Fab, 7 of which exhibited class 4 and 5 epitope competition patterns. The latter VHH-Fcs recognize an epitope at the class 4/class 5 interface that was described previously for anti-RBD VHHs.^61,62^

We also evaluated the potencies of the VHH-Fcs as assessed by inhibitory concentration at 50% values (IC50s) in a pseudovirus neutralization assay^63^ (Figure 4C). Class 4 VHH-Fcs exhibited the greatest neutralization breadth in our panel with neutralizing titers ≤2 µg/mL for 6 of 9 strains tested, including 3 SARS-2 VOCs. Class 4/5 VHH-Fcs exhibited less neutralizing breadth, with titers ≤2 µg/mL against 5 of 9 strains (P2-A12), 4 of 9 strains (P1-B5), 3 of 9 strains (P1-F6, P1-G11, P1-H1, P2-C12), or 2 of 9 strains (P2-D6). All class 4/5 VHH-Fcs exhibited neutralizing activity against KP.3, the most recent VOC we evaluated, whereas class 4 VHH-Fcs were less potent against KP.3.

### VHHs recognize conserved RBD epitopes

To further characterize VHH recognition, we solved crystal structures of selected VHHs complexed with SARS-2 RBD (Table S3). A 1.8 Å resolution P1-F6–RBD complex structure revealed VHH interactions involving complementarity determining regions 1, 2, and 3 (CDR1, CDR2, and CDR3) and framework region 3 (FWR3) (Figure 5A; Figure S4). A 1.5 Å resolution structure of a P1-G11–RBD complex revealed a similar angle of approach for RBD recognition by the P1-G11 VHH as found for the P1-F6 VHH (Figure 5B), and like P1-F6, P1-G11 engaged RBD through its three CDRs plus FWR3 (Figure 5B). Despite sharing only 85% amino acid sequence identity in the VHH domain (Figure 4D), the binding poses and epitopes of P1-F6 and P1-G11 were similar, comprising conserved residues in the class 4 and class 5 RBD epitopes, consistent with competition of these VHHs with class 1/4, 4, and 5 Fabs (Figure 4B) and broad binding to RBDs from different sarbecovirus clades (Figure 4A). Based on binding, competition, and structure results, we classify the P1-F6 and P1-G11 VHHs as class 4/5 anti-RBD mAbs.

**Figure 5.**
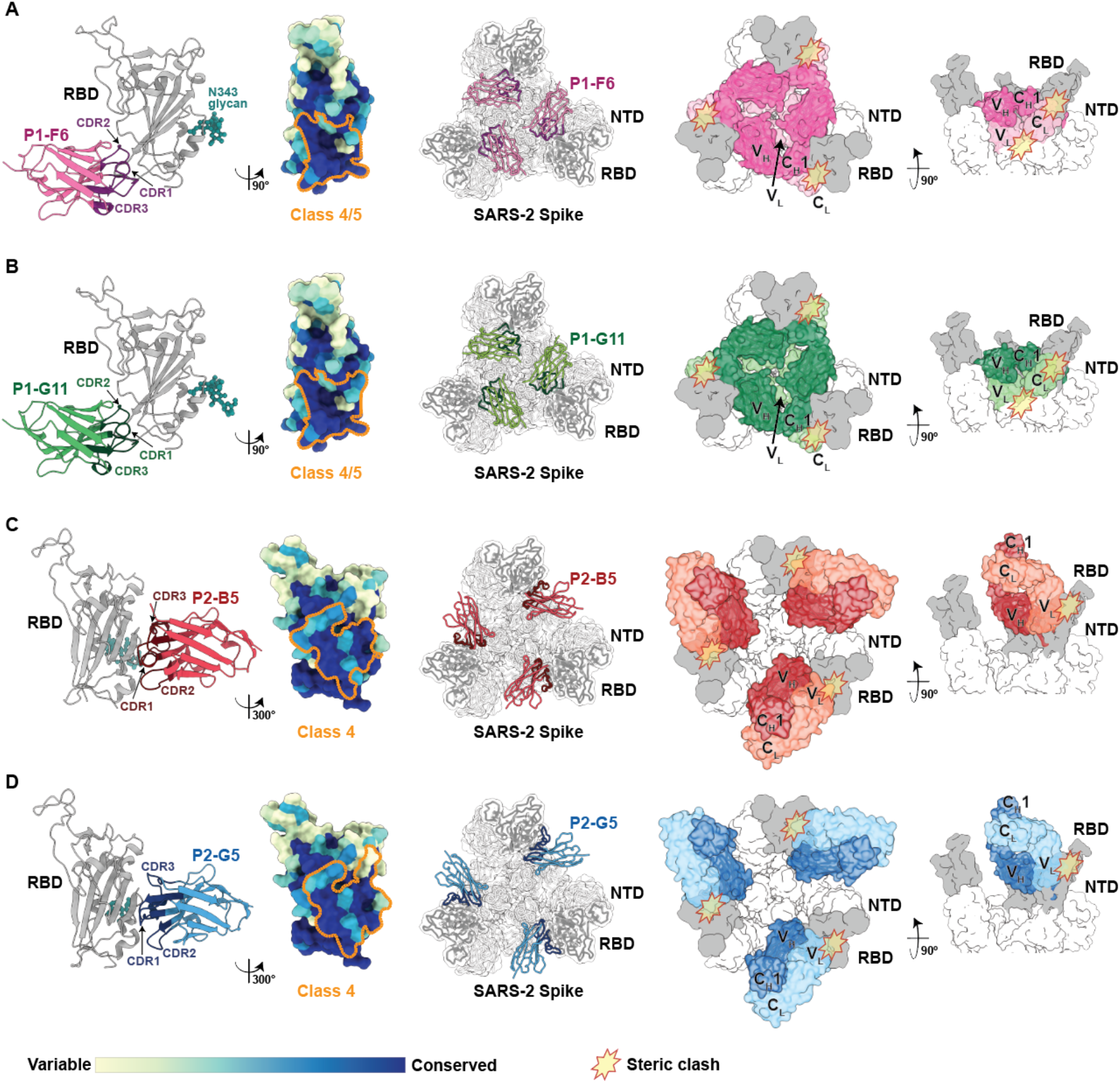
VHH-RBD structures demonstrate recognition of conserved RBD epitopes that are inaccessible to analogous VL-containing two domain Fabs. Five panels related to structures of RBD complexes with (A) P1-F6, (B) P1-G11, (C) P2-B5, and (D) P2-G5 VHHs are shown. Left: Cartoon diagram of the VHH-RBD structure with RBD in gray and the N343 N-glycan in teal, and VHH (pink, green, red, or cyan) with CDRs highlighted in a darker color. Second from left: Epitope for each VHH outlined in orange dots on an RBD surface colored to distinguish variable and conserved RBD regions (calculated from 16 sarbecovirus sequences using ConSurf^74^). Middle: Top-down view of a VHH-spike trimer complex modeled using the VHH-RBD structure and SARS-2 spike coordinates from PDB: 7RKV. Right panels: Top view (second from right) and side view (right) of a model of a SARS-2 spike trimer surface complexed with a four-domain Fab (VH, VL, CH1, CL) derived by superimposing the coordinates of the VH domain of the human mAb C118 (PDB: 7RKV) onto the VHH coordinates of the VHH-spike trimer model in the middle panel. Predicted steric clashes between a spike trimer and a Fab containing a VH domain with an RBD-recognizing pose analogous to a VHH are indicated by yellow starbursts.

The structures of RBD complexes with P2-B5 and P2-G5 VHHs (1.5 Å and 1.9 Å resolution, respectively), which share only 70% sequence identity (Figure 4D), demonstrated recognition of the conserved class 4 RBD epitope using similar approach angles (Figure 5C, D), consistent with binding and competition results (Figure 4). Both VHHs utilized all three CDRs for interactions with the RBD (Figure 5C, D; Figure S4, S6A, B). Notably, these VHHs used CDR3 mainchain atoms to form hydrogen bonds with mainchain atoms in an RBD β-strand (residues 375-381) to extend an RBD β-sheet (Figure S6A,B). P2-B5 also stabilized its interaction with RBD using an additional hydrogen bond between a CDR3 sidechain and the N-terminus of the RBD β-strand, and P2-G5 included a stabilizing sidechain contact between a CDR1 residue and the RBD β-strand C-terminus (Figure S6A,B). Previous studies of human mAb-RBD interactions also described backbone hydrogen bonding interactions between heavy chain CDR3s (CDRH3s) that included a YYDxxG motif and RBD residues 375-381, thereby extending the RBD β-sheet,^64^ and we observed analogous interactions with RBDs for mAbs elicited in mosaic-8b–immunized rabbits, although the rabbit Abs contained only the YY portion of the YYDxxG motif.^23^ Here, we show that the RBD recognition strategy involving extension of the RBD β-sheet can be accomplished by llama VHHs with SYLHYY (P2-B5) and SWLLRV (P2-G5) CDR3 sequences rather than a YYDxxG motif (Figure S6). As another example, the crystal structure of a SARS-1 RBD complex with VHH-72, a llama anti-RBD VHH raised against the SARS-1 spike,^65^ also showed mainchain hydrogen bonds between CDR3 residues and an RBD β-strand to extend the RBD β-sheet (Figure S6C). These results illustrate distinct mechanisms by which VH-VL and VHHs from different organisms (humans, rabbits, and llamas) arrive at a common mode of RBD recognition that involves mainchain hydrogen bonds to extend an RBD β-sheet.

We also used our VHH-RBD structures to address whether a two-domain VHVL-containing Fab could access the epitope recognized by the corresponding VHH in the context of a spike trimer. Starting with the spike trimer coordinates from a Fab-spike complex, we modeled the interactions of each VHH with the RBDs in a spike trimer (Figure 5, middle panels), finding that the structurally-characterized class 4/5 and class 4 VHHs described above can only be accommodated on an RBD in an “up” position, similar to Fab-containing Abs recognizing cryptic RBD epitopes.^17^ Modeling of VHHs onto the RBDs of a spike trimer structure suggested that both class 4/5 VHHs (P1-F6 and P1-G11) can recognize “up” RBDs with no predicted steric clashes with nearby regions on the spike and the VHHs targeting RBD regions in close proximity to the spike S2 subunit (Figure 5A, B). In addition, modeling predicted that the class 4 VHHs (P2-B5 and P2-G5) interacted with upper regions of “up” RBDs with no predicted steric clashes in the context of a spike trimer (Figure 5C, D). In all modeled VHH-spike structures, three VHHs could be accommodated on a spike trimer (Figure 5, middle panels). We then asked whether a larger Fab-containing Ab whose VH domain interacts with RBDs with the same approach angle as these VHHs could be accommodated on a spike trimer. For this modeling, we superimposed the VH domain of a Fab structure on the VHH portion of a VHH-spike trimer model (right panels in Figure 5) and assessed whether any of the remaining domains of the Fab (VL, CH1, CL) were predicted to clash with parts of the spike trimer. For each of the Fab-spike models, we found predicted clashes with one or more of the other Fab domains (two right panels in Figure 5). These results are consistent with the hypothesis that VHHs can target epitopes that cannot accommodate Abs with light chain-including Fabs.

VHH-VL models were also used to visualize the locations of CDR3s with respect to the VHH surface that would interact with the VL domain in a conventional Fab (Figure S7). For each of the VHHs for which we derived VHH-RBD structures, the CDR3s included a portion that was folded back to interact with the side of the VHH that would associate with a partner VL domain in a two-domain Fab. For our VHH-RBD structures, the folded back CDR3s resulted in an often observed “sideways” interaction of a VHH with antigen; e.g., as seen in the VHH-72–RBD complex structure^65^ (Figure S6C) and other camelid VHH-antigen interactions.^66^ A sideways orientation of a VHH with respect to antigen can occur when a long CDR3 loop interacts with the side of the VHH that would interact with the VL domain in a conventional Fab,^66^ consistent with the relatively long CDR3s in P1-F6, P1-G11, P2-G5, and P2-B5 (19, 19, 22, and 19 amino acids, respectively) compared with the average CDRH3 length in human IgGs (15.5 ± 3.2 residues).^67^ This can result in an angle of approach that is favorable for recognition of more occluded epitopes.

## Discussion

Heavy chain-only VHH nanobodies offer advantages over conventional heavy chain-light chain mAbs. For example, the smaller size of VHH domains compared with Fabs allows greater access to occluded epitopes and more efficient tissue penetration, and VHH nanobodies are highly soluble and stable across broad temperature and pH conditions.^30,31^ For these reasons, nanobodies are increasingly being developed and used as therapeutics, with 30-50 VHH-based candidates in clinical trials in the US.^68^ Nanobodies recognizing conserved regions of sarbecovirus spike proteins would be a valuable addition to therapeutics for combatting continued threats from new variants of SARS-2 and potential zoonotic sarbecovirus spillovers. Of relevance for this goal, structural studies of >1,100 anti-SARS-2 Abs revealed enrichment of conserved and cryptic epitopes for nanobodies as compared with conventional Fab-containing Abs.^62^ Thus, the intrinsic structural features of nanobodies make them well suited for targeting functionally constrained epitopes of RBD that are difficult for conventional Abs to access.

Nanobodies are typically identified using synthetic VHH display libraries or from immunized camelids such as llamas or alpacas.^30,31^ Although synthetic libraries offer a powerful and readily accessible platform for nanobody discovery, immunization allows sampling of naturally elicited VHH diversity and affinity-matured responses. Many commonly used synthetic VHH libraries employ relatively fixed FWRs as a constant scaffold while varying CDRs within discrete limits such as binned CDR lengths.^69,70^ For example, an often-used yeast display nanobody library samples CDR3 length diversity at three discrete values: 7, 11, and 15 amino acids.^69^ In contrast, immunization allows exploration of CDR and FWR space simultaneously, as well as subjecting these combinations to *in vivo* B cell activation and selection. Notably, we observed FWR contacts with RBD in the four VHH-RBD solved in this study (Figure S4), residues that would not typically be included in VHH libraries. Thus, immunization can co-select binding geometry and chemistry in ways that are difficult to recapitulate using a fixed synthetic scaffold.

Here, we exploited the natural VHH repertoire by immunizing a llama with mosaic-8b RBD-NP, a pan-sarbecovirus vaccine candidate that was shown to preferentially elicit broadly cross-reactive mAbs in animals that produce conventional Abs.^22,23^ We found that immunization of a llama with mosaic-8b NP resulted in elicitation of VHH Abs against conserved RBD regions (e.g., class 4, 1/4, 5, and 4/5). These conserved epitopes are optimal targets for therapeutic Abs, as compared to variable class 1 and class 2 RBD epitopes recognized by conventional mAbs and VHHs elicited by RBD and spike immunogens.^21,24–26,60,62,71^ This finding extends previous mAb discovery studies using mosaic-8 RBD-NP immunogens^22,23^ to the camelid nanobody repertoire. Our results suggest that the combination of mosaic-8b RBD-NPs and the distinctive structural diversity of the camelid VHH repertoire is an effective strategy to enrich nanobodies against conserved RBD epitopes. Importantly, the VHHs recovered in this study encompassed multiple CDR3 lengths and contained RBD contact residues from both CDRs and FWRs, illustrating repertoire diversity that would not be fully recapitulated using synthetic libraries in which FWR sequences and CDR lengths are constrained.

Use of the Beacon optofluidics system enabled characterization of llama memory B cells using sequential multiplexed assays to create a detailed profile of IgGs elicited following immunization of a llama (Figure 2; Figure S2, S3). Our initial B cell enrichment strategy prioritized antigen binding, with an IgG2/3 pulldown as a secondary enrichment. We observed broad antigen binding reactivity in the full dataset; however, IgG1+ B cells that do not encode VHHs were also present (Figure 3B). In an independent experiment, we reversed the enrichment order by prioritizing IgG2/3 pulldown before antigen binding enrichment, which provided excellent enrichment for IgG2 but reduced frequency of antigen-reactive B cells. We therefore elected to use both antigen reactivity and IgG2 signal to prioritize the selection of cells for export and subsequently relied on sequence information for definitive subclass determination. This strategy avoids requiring either antigen binding or subclass enrichment alone to serve as an absolute selection criterion, offering flexibility during discovery.

Sequence recovery was high for exported cells throughout our cloning pipeline, which incorporated both an initial PCR product sequencing approach to determine the IgG subclass and recover the VHH sequence, followed by cloning the VHH into a Fc-fusion expression plasmid and sequence verification. To our knowledge, our study is the first to report using a Beacon and commercially available media reagents for isolation, culture, and characterization of B cells secreting HC-IgGs from immunized llamas, as opposed to a previously published study in which proprietary media was used to characterize heavy chain-only Abs from alpacas using a Beacon instrument.^72^ The use of readily accessible reagents could facilitate adoption and adaptation of this workflow for VHH discovery against other pathogens and therapeutic targets.

An important advantage of the Beacon-based approach was the ability to evaluate individual B cells through a serial multiplexed assay. We evaluated recognition of 15 individual sarbecovirus RBDs in addition to IgG subclass determination (Figure 3A; Figures S2A, S3). This allowed cross-reactivity to be assessed directly at the antibody-secreting B cell stage rather than relying on sequence recovery followed by recombinant Abs expression and retrospective characterization. Moreover, this strategy also allows B cells exhibiting desirable binding profiles to be prioritized for export and cloning, enabling discovery of VHHs with favorable binding and neutralization properties that could be further developed as therapeutics (Figure 4).

Clinical experience with other VHH-based molecules also supports the feasibility of this therapeutic modality. Of particular relevance to potential therapeutic use in humans, a phase 1 clinical trial showed that XVR011-Fc (an affinity matured version of VHH-72^37^) was well tolerated and exhibited favorable pharmacokinetic and immunogenicity profiles in both healthy and SARS-2–infected individuals.^37^ However, this trial was prematurely discontinued in 2022 because XVR011 did not exhibit potency against the SARS-2 Omicron BA.2 variant, which was emergent at that time. Notably, XVR011 exhibited reduced binding breadth compared to the VHH-Fcs isolated in our study (Figure 4A). The loss of activity of XVR011 against recent variants does not diminish the potential utility of the VHH platform; rather, it highlights the importance of selecting VHHs that recognize conserved epitopes and demonstrate breadth against emerging SARS-2 variants and zoonotic sarbecoviruses.

In summary, our results demonstrate that llama immunization with mosaic-8b RBD-NPs coupled with optofluidic, multiplex individual B-cell screening provides an efficient strategy to identify naturally elicited VHHs directed against conserved sarbecovirus RBD epitopes. This work also illustrates how rational immunogen design can be integrated with the natural diversity of camelid VHH repertoires and high-throughput single cell screening to discover Abs that may be difficult to obtain from conventional Ab responses or synthetic libraries. Given favorable clinical profiles with VHH-Fcs^68^ and the continuing evolution of SARS-2 and potential zoonotic sarbecovirus spillovers,^73^ the broadly reactive VHHs identified in this study warrant further development as potential prophylactic and therapeutic agents against current SARS-2 VOCs and future sarbecovirus spillovers.

## Acknowledgments

We thank Ian Fotheringham, Leonardo Magneschi, Scott Baxter, Rita Cruz, Martina Quintanar-Audelo, Ellis Robb, and Cristina Serrano-Amatriain at Ingenza, Ltd. for providing mosaic-8b RBD-NPs, Gary Stephens, Barney Jones, Hong Lin, and Basil (a llama) at the University of Reading for llama immunizations, Jost Vielmetter (Caltech Beckman Institute Protein Expression Center) and Diana Melnyk (MRC Weatherall Institute of Molecular Medicine, Oxford University) for protein production, the Caltech Beckman Institute Beacon Center for Single Cell Biology for conducting Beacon experiments, and Marta Murphy for assistance with figures. We also thank Jens Kaiser, staff at Stanford Synchrotron Radiation Lightsource, and the Caltech Molecular Observatory for X-ray data collection support. Use of the Stanford Synchrotron Radiation Lightsource, SLAC National Accelerator Laboratory, is supported by the U.S. Department of Energy, Office of Science, Office of Basic Energy Sciences under Contract No. DE-AC02-76SF00515. The SSRL Structural Molecular Biology Program is supported by the DOE Office of Biological and Environmental Research, and by the National Institutes of Health, National Institute of General Medical Sciences (including P30GM133894). These studies were funded by the Caltech Merkin Institute (P.J.B.), Wellcome Leap (P.J.B.), and the Beckman Institute at Caltech (K.E.M.).

## Competing Interest Statement

P.J.B. is an inventor on a US patent application (17/523,813) filed by the California Institute of Technology that covers mosaic RBD-NPs.

