## Supplemental Figures for "A mosaic RBD-nanoparticle pan-sarbecovirus vaccine elicits nanobodies that target highly conserved epitopes"

### Supplementary Figures

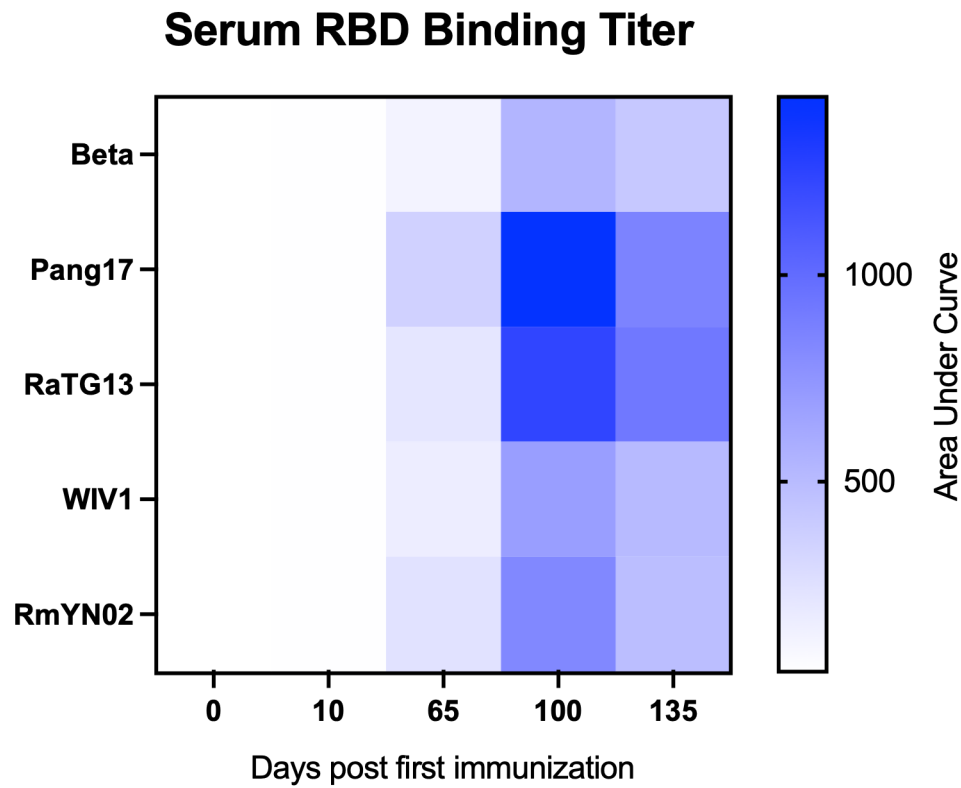

Figure S1: Serum Ab responses post mosaic-8b immunization. Serum Ab binding titer against RBDs (Beta, Pang17, RaTG13, WIV1, or RmYN02) at various timepoints post immunization presented as a heat map. Each data point represents the mean of two AUC values, with color intensity corresponding to the magnitude of the serum Ab response against the indicated RBD.

**A**

| Acquisition channel | Assay 1 | Assay 2 | Assay 3 | Assay 4 | Assay 5 |
| --- | --- | --- | --- | --- | --- |
| FITC | Anti-Alpaca IgG2/3 | Anti-Alpaca IgG2/3 | Anti-Alpaca IgG2/3 | XBB1.5 RBD (clade 1b) | Goat Anti-Alpaca IgG2/3, Fab |
| TRED | Khоста-2 RBD (clade 3) | SARS-2 Beta RBD (clade 1b) |  | Pang17 RBD (clade 1b) | WIV1 RBD (clade 1a) |
| Cy5 | BtkY72 RBD (clade 3) | Anti Alpaca IgG (H+L) | Anti Alpaca IgG (H+L) | Goat Anti-Alpaca VHH, Fab | SHC014 RBD (clade 1a) |
| PE | Yun11 RBD (clade 2) | BM48-31 RBD (clade 3) |  | RaTG13 RBD (clade 1b) | Rs4081 RBD (clade 2) |
| DAPI | SARS-1 RBD (clade 1a) | SARS-2 KP.3 RBD (clade 1b) |  | RmYN02 RBD (clade 2) | Rf1 RBD (clade 2) |
| bead type | Anti-camelid IgG (H+L) | Anti-camelid IgG (H+L) | SpyCatcher003-mi3 | Protein G | Protein G |

Matched for mosaic-8b

**B**

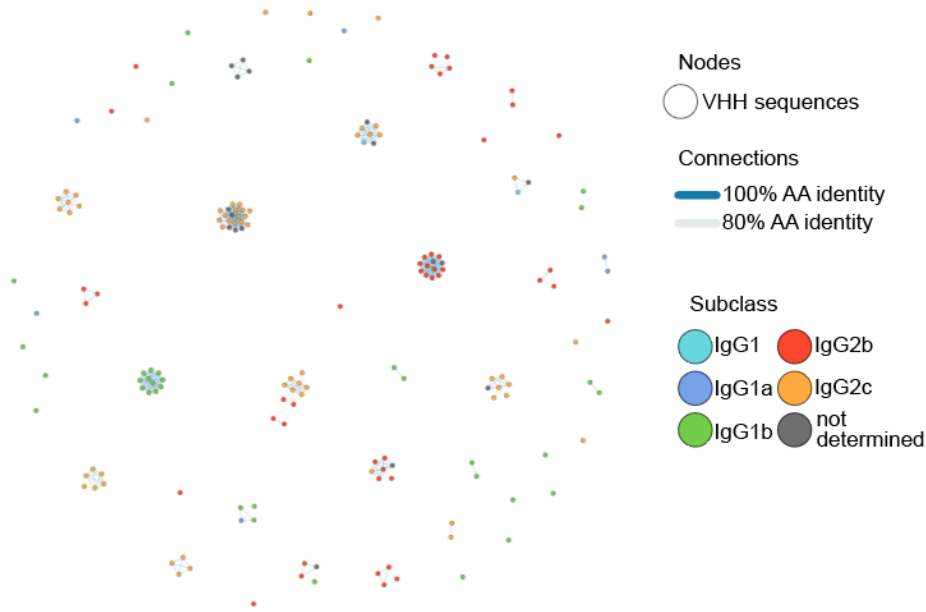

Figure S2. Assay and sequencing results for llama memory B cells isolated and characterized on the Beacon. (A) Serial multiplexed assay strategy used to characterize llama memory B cells. Each assay used a capture bead and fluorescently labeled RBDs and secondary Abs to visualize the subclass of secreted Abs and their RBD-binding properties as they accumulate on the stationary capture beads. RBD clades are indicated after the RBD name, and the fluorophore used for labeling is indicated in the Acquisition channel column. RBDs other than those that were matched to the mosaic-8b RBD-NP immunogen (SARS-2 Beta, RaTG13, Pang17, WIV1, SHC014, Rs4081, RmYN02, Rf1) are mismatched. SpyCatcher003-mi3<sup>35</sup> is the protein NP scaffold used to generate the mosaic-8b RBD-NP immunogen. Signal was measured above the nanopen compartment in which the cell secreting these Abs had been placed in culture within the microfluidic chip. (B) Cluster analysis of all recovered VH sequences (IgG1 and IgG2) represented by nodes colored by subclass. Cluster and connections between nodes represent sequence relationships among recovered VH sequences.

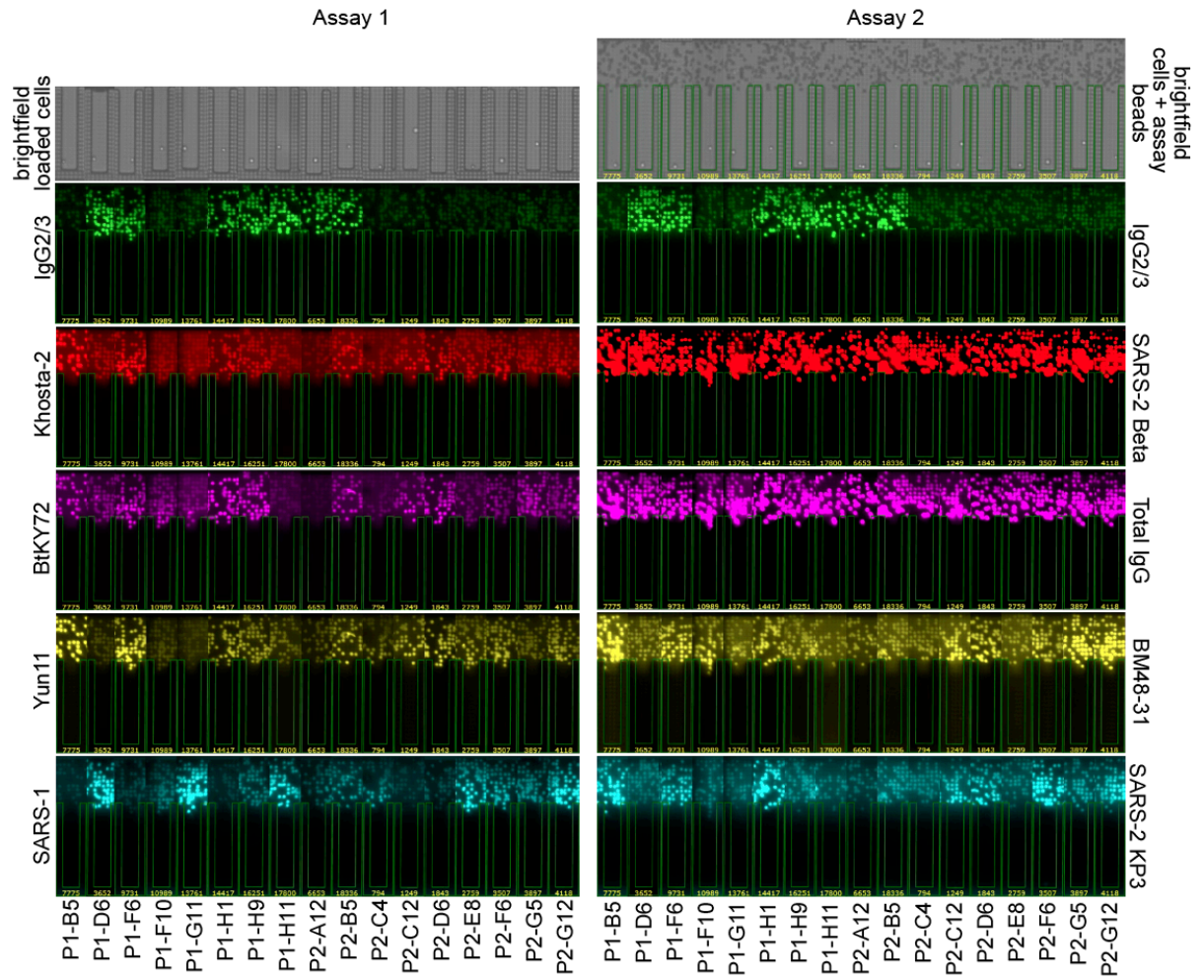

Figure S3. Images of individual nanopen assay 1 and 2 (Figure S2) results for cells of interest that were exported, VHH sequences cloned, and for which characterization is reported.

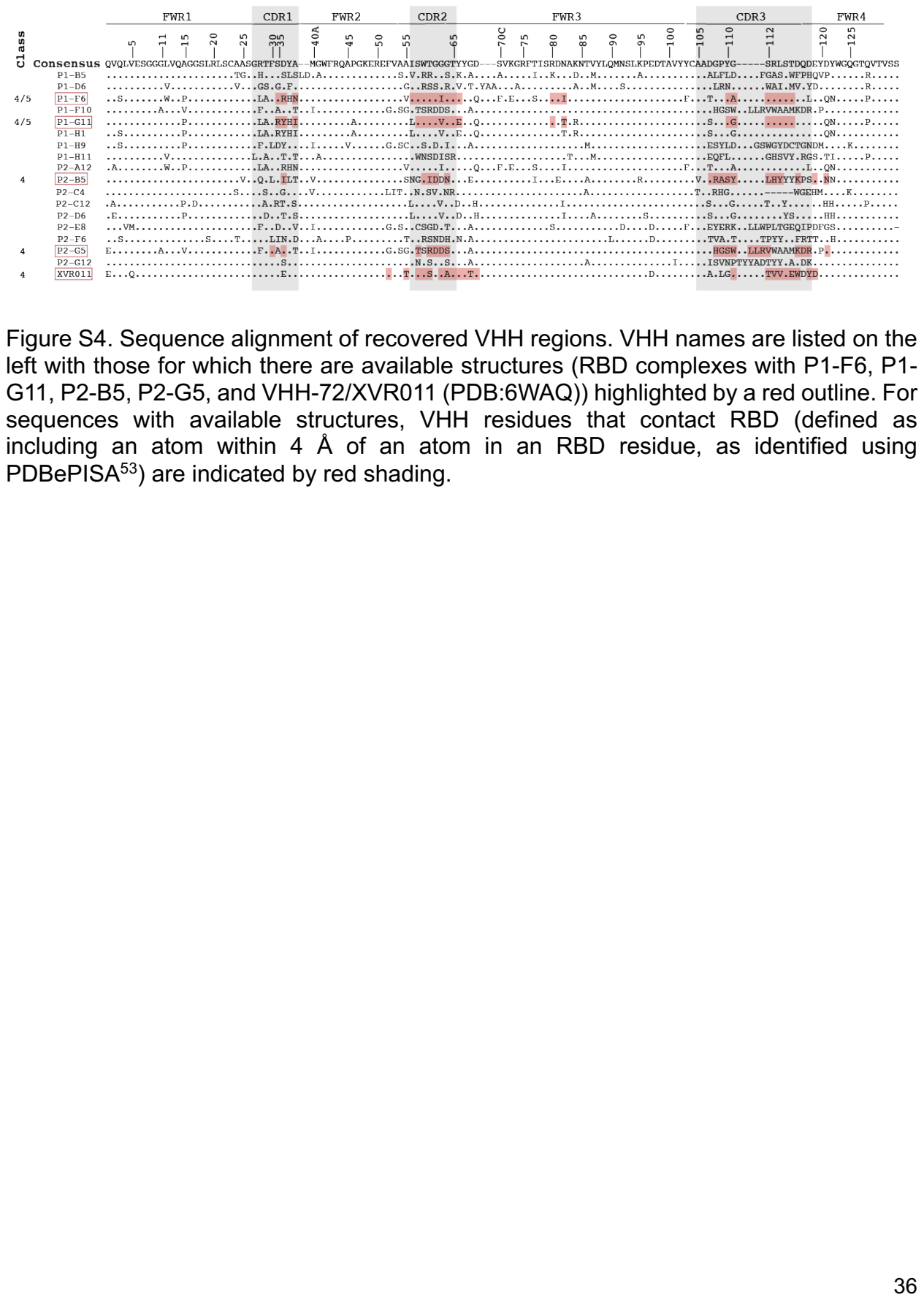

Figure S4. Sequence alignment of recovered VHH regions. VHH names are listed on the left with those for which there are available structures (RBD complexes with P1-F6, P1- G11, P2-B5, P2-G5, and VHH-72/XVR011 (PDB:6WAQ)) highlighted by a red outline. For sequences with available structures, VHH residues that contact RBD (defined as including an atom within 4 Å of an atom in an RBD residue, as identified using PDBePISA53) are indicated by red shading.

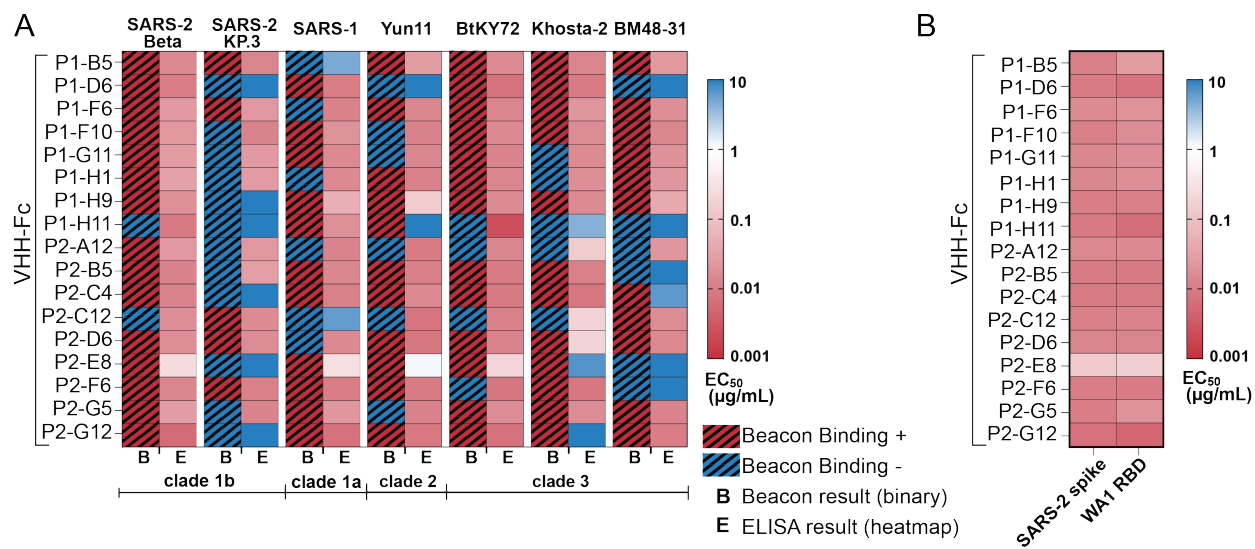

Figure S5. Beacon and ELISA binding results. (A) Comparison of binding results from Beacon multiplexed assays (labeled as B across the bottom) and ELISAs (labeled as E across the bottom). Beacon results are shown as positive for binding (red crosshatched squares) or negative for binding (blue crosshatched squares). ELISA results are shown in squares as a gradient depending on EC<sub>50</sub> value. (B) ELISA comparison of VHH-Fcs binding to the SARS-2 WA1 spike trimer or the SARS-2 WA1 RBD.

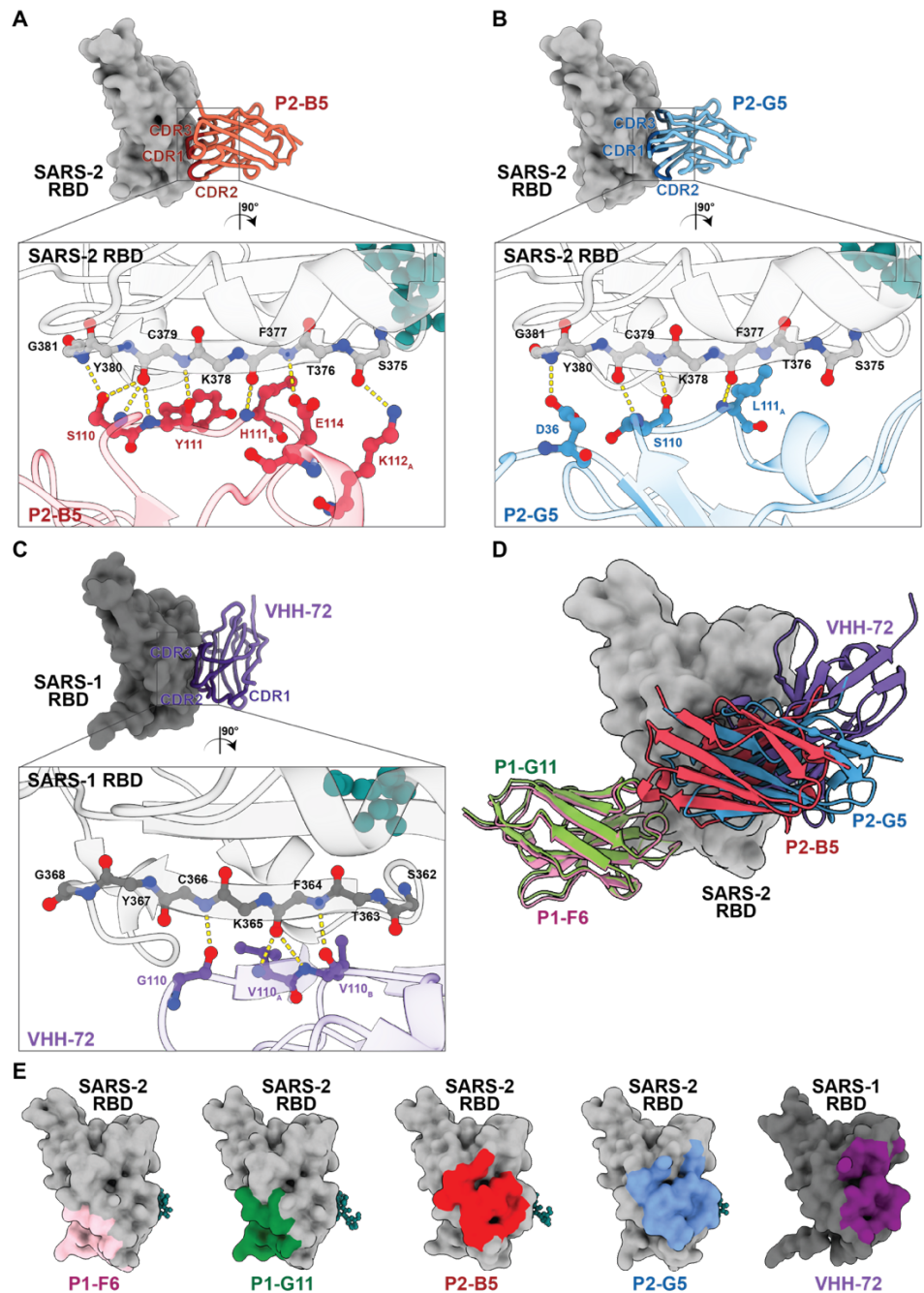

Figure S6. Examples of VHH-RBD interfaces that include mainchain hydrogen bonding interactions. (A-C) Top: Each structure is shown with the RBD as a grey surface and the VHH as a carbon-α trace with CDRs and FWR3 in darker colors. Bottom: Zoomed-in views showing hydrogen bonds at VHH-RBD interfaces (yellow dotted lines) between atoms in amino acid residues as ball-and-stick representations (main chain oxygen atoms in red; main chain nitrogen atoms in blue; N343 N-glycan in teal). (D) RBD surface with overlay of four VHHs from this study plus VHH-72<sup>65</sup> (PDB:6WAQ), a VHH raised against SARS-1 RBD that was affinity matured to create XVR011,<sup>37</sup> a VHH-Fc evaluated in a clinical trial.<sup>75</sup> (E) Comparison of epitopes for VHHs on RBD surfaces.

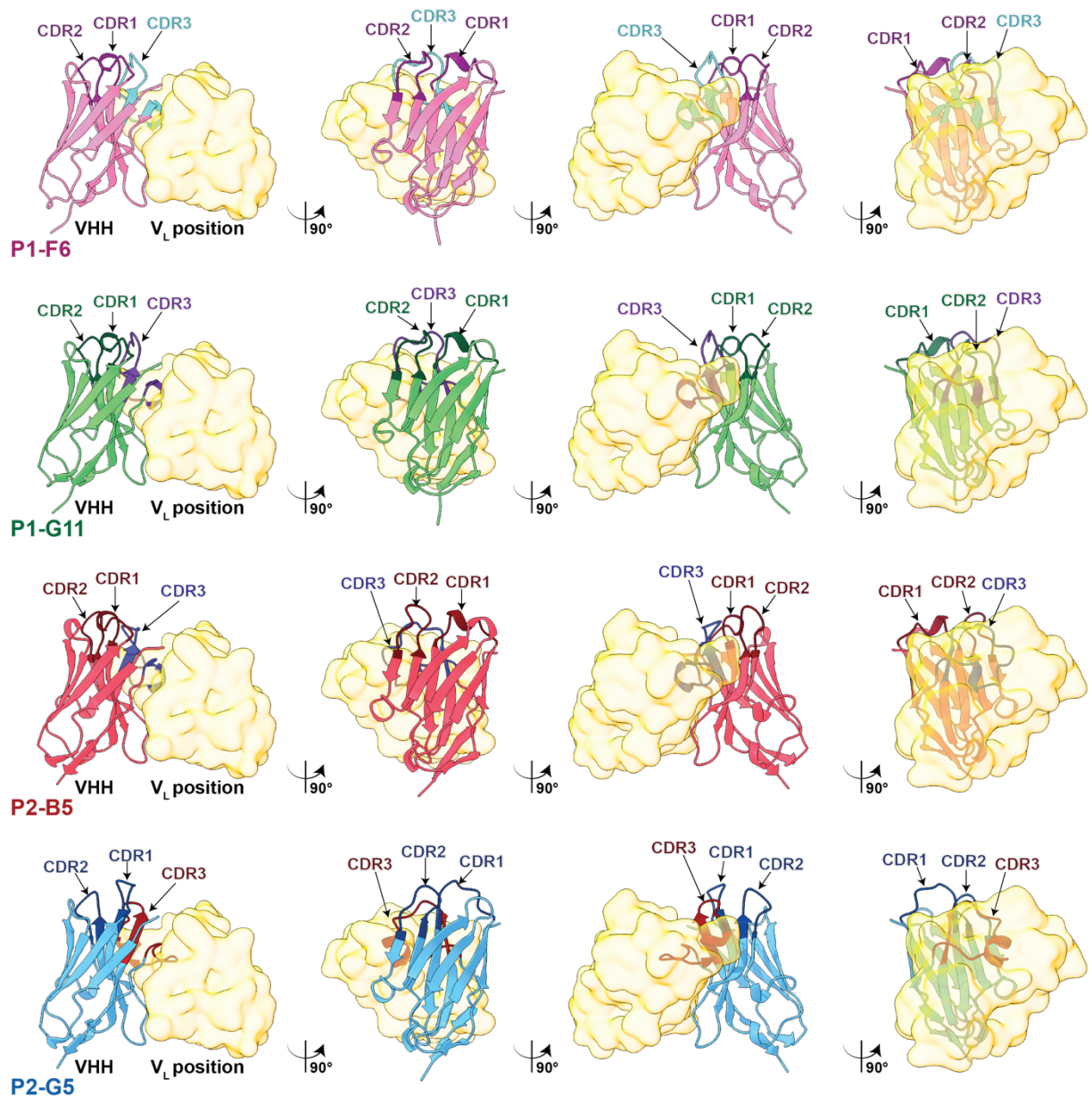

Figure S7. CDR3s of VHs from VHH-RBD structures adopt a bent conformation. VHs are shown in cartoon representation with CDRs in darker colors (CDRs 1 and 2) or a contrasting color (CDR3). To demonstrate folding back of the VHH CDR3 into the region that would normally be occupied by a V<sub>L</sub> domain in a conventional Fab, the V<sub>L</sub> domain location is shown as a yellow surface derived from a model of a four-domain Fab (V<sub>H</sub>, V<sub>L</sub>, C<sub>H</sub>1, C<sub>L</sub>) made by superimposing the coordinates of the V<sub>H</sub> domain of the human mAb C118 (PDB: 7RKS) onto the VHH coordinates.

| Name | Sequence |
| --- | --- |
| library forward (TrueSeqRead1) | TCGTCGGCAGCGTCAGATGTGTATAAGAGACAGTACACGACGCTCTTCCGATCT |
| camelid hinge-CH2 reverse | GTCTCGTGGGCTCGGAGATGTGTATAAGAGACAGRCATTTGGGACACTTGGA |
| VHH F1 | CCACTTGCTCTTCAAGCCAGGTGCAGCTGGTRGAGTC |
| VHH F2 | CCACTTGCTCTTCAAGCCAGGCGCAGCTGGTGGAGTC |
| VHH F3 | CCACTTGCTCTTCAAGCCAGGAGCAGCTGGTAGAGTC |
| VHH F4 | CCACTTGCTCTTCAAGCCAGGTGCAGCTGGTAGAGTC |
| VHH F5 | CCACTTGCTCTTCAAGCCAGGTGTCGCTGGTAGAGTC |
| VHH F6 | CCACTTGCTCTTCAAGCCAGGTGCAGCTCGTGGAG |
| VHH F7 | CCACTTGCTCTTCAAGCCAAGTGGACCTGATAGAGTC |
| VHH F8 | CCACTTGCTCTTCAAGCCAGGTGGCCCTAATAGAGTC |
| VHH F9 | CCACTTGCTCTTCAAGCCAGGTGCAGGTGATGGAGTC |
| VHH R1 | GGGATTGCTCTTCACGATGAGGAGACGGTGACCTGGG |
| VHH R2 | GGGATTGCTCTTCACGAAGAGGAGACGGTGACCTGGG |
| VHH R3 | GGGATTGCTCTTCACGACGAGGAGACGGTGACCGGGG |
| VHH R4 | GGGATTGCTCTTCACGATGAGGAGACGGTGACCCGGG |

Table S1. Primers used to amplify VHH regions from cDNA library for IgG subclass determination and to amplify VHH regions for cloning into an expression vector including a gene encoding human IgG1 Fc.

| **VHH** | **V-GENE** | **J-GENE** | **CDR1**<br>**Length** | **CDR2**<br>**Length** | **CDR3**<br>**Length** | **subclass** |
| --- | --- | --- | --- | --- | --- | --- |
| P1-B5 | IGHV1S1 | IGHJ4 | 8 | 8 | 20 | IgG2b |
| P1-D6 | IGHV1S2 | IGHJ4 | 8 | 8 | 18 | IgG2c |
| P1-E6 | IGHV1S2 | IGHJ4 | 8 | 8 | 19 | IgG2c |
| P1-F6 | IGHV1S1 | IGHJ4 | 8 | 8 | 19 | IgG2c |
| P1-F10 | IGHV1S1 | IGHJ4 | 8 | 8 | 22 | IgG2b |
| P1-G11 | IGHV1S2 | IGHJ4 | 8 | 8 | 19 | IgG2c |
| P1-G12 | IGHV1S1 | IGHJ4 | 8 | 8 | 19 | IgG2c |
| P1-H1 | IGHV1S2 | IGHJ4 | 8 | 8 | 19 | IgG2c |
| P1-H9 | IGHV1S1 | IGHJ4 | 8 | 8 | 21 | IgG2c |
| P1-H11 | IGHV1S4 | IGHJ4 | 8 | 8 | 18 | IgG2c |
| P2-A12 | IGHV1S2 | IGHJ4 | 8 | 8 | 19 | IgG2c |
| P2-B5 | IGHV1S3 | IGHJ4 | 8 | 8 | 19 | IgG2c |
| P2-C4 | IGHV1S4 | IGHJ4 | 8 | 8 | 13 | IgG2b |
| P2-C12 | IGHV1S4 | IGHJ4 | 8 | 8 | 19 | IgG2b |
| P2-D6 | IGHV1S2 | IGHJ4 | 8 | 8 | 19 | IgG2b |
| P2-E8 | IGHV1S1 | IGHJ6 | 8 | 8 | 22 | IgG2b |
| P2-F6 | IGHV1S5 | IGHJ4 | 8 | 8 | 20 | IgG2b |
| P2-G5 | IGHV1S1 | IGHJ4 | 8 | 8 | 22 | IgG2b |
| P2-G12 | IGHV1S4 | IGHJ4 | 8 | 8 | 24 | IgG2b |
| XVR011 | IGHV1S2 | IGHJ4 | 8 | 8 | 18 | Not<br>determined |

Table S2. Sequence features of VHHs described in this study.

| Structures | P1-F6 VHH-RBD | P1-G11 VHH-RBD | P2-B5 VHH-RBD | P2-G5 VHH-RBD |
| --- | --- | --- | --- | --- |
| Wavelength (Å) | 0.9795 | 0.9795 | 0.9795 | 0.9795 |
| Resolution range (Å) | 37.68 - 1.8 (1.84 - 1.8) | 39.06 - 1.5 (1.54 - 1.5) | 38.22 - 1.5 (1.54 - 1.5) | 39.61 - 1.9 (1.95 - 1.9) |
| Space group | P 2<sub>1</sub> 2<sub>1</sub> 2<sub>1</sub> | P 2<sub>1</sub> 2<sub>1</sub> 2<sub>1</sub> | C 2 | P 2<sub>1</sub> 2<sub>1</sub> 2<sub>1</sub> |
| Unit cell (Å, °) | 46.3 64.8 123.6 90 90 90 | 47.5 68.5 123.0 90 90 90 | 128.1 55.2 62.5 90 97.6 90 | 58.2 71.4 108.1 90 90 90 |
| Total reflections (no.) | 475,753 (33,518) | 878,890 (62,505) | 478,713 (33,976) | 475,255 (22,490) |
| Unique reflections (no.) | 35,213 (2,457) | 64770 (4,542) | 68,645 (4,893) | 36,137 (2,433) |
| Multiplicity | 13.5 (13.6) | 13.6 (13.8) | 7.0 (6.9) | 13.2 (9.2) |
| Completeness (%) | 99.6 (99.1) | 99.3 (98.5) | 98.8 (96.2) | 99.3 (93.1) |
| Mean I/sigma(I) | 12.4 (1.3) | 10.85 (1.2) | 12.44 (1.4) | 10.8 (0.8) |
| Wilson B-factor (Å<sup>2</sup>) | 25 | 22 | 20 | 30 |
| R-merge | 0.132 (1.826) | 0.112 (1.434) | 0.070 (1.028) | 0.145 (2.123) |
| R-meas | 0.137 (1.896) | 0.117 (1.488) | 0.075 (1.111) | 0.150 (2.248) |
| R-pim | 0.037 (0.506) | 0.032 (0.394) | 0.028 (0.417) | 0.041 (0.721) |
| CC1/2 | 0.999 (0.596) | 0.997 (0.786) | 0.998 (0.701) | 0.999 (0.615) |
| CC* | 1 (0.864) | 0.999 (0.938) | 1 (0.908) | 1 (0.873) |
| Reflections used in refinement (no.) | 35,209 (2,457) | 64,759 (4,538) | 68,672 (4,761) | 36,043 (2,395) |
| Reflections used for R-free (no.) | 2,000 (140) | 2,000 (140) | 1,998 (138) | 1,999 (132) |
| R-work | 0.170 (0.228) | 0.158 (0.237) | 0.151 (0.219) | 0.173 (0.337) |
| R-free | 0.213 (0.299) | 0.1928 (0.302) | 0.181 (0.243) | 0.223 (0.443) |
| Number of non-hydrogen atoms | 2,827 | 2,972 | 3,128 | 2,903 |
| Macromolecules | 2,562 | 2,601 | 2,693 | 2,615 |
| Ligands | 44 | 83 | 78 | 94 |
| Solvent | 221 | 288 | 357 | 194 |
| Protein residues (no.) | 322 | 322 | 321 | 325 |
| RMS (bonds) (Å) | 0.008 | 0.005 | 0.005 | 0.008 |
| RMS (angles) (°) | 0.89 | 0.73 | 0.82 | 0.85 |
| Ramachandran favored (%) | 98.4 | 98.1 | 98.7 | 98.4 |
| Ramachandran allowed (%) | 1.6 | 1.9 | 1.3 | 1.6 |
| Ramachandran outliers (%) | 0 | 0 | 0 | 0 |
| Rotamer outliers (%) | 0.4 | 0 | 0.6 | 0.7 |
| Clashscore | 1.6 | 1.1 | 3.5 | 0.8 |
| Average B-factor (Å<sup>2</sup>) | 32 | 30 | 27 | 41 |
| Macromolecules | 31 | 28 | 25 | 40 |
| Ligands | 72 | 62 | 51 | 65 |
| Solvent | 35 | 38 | 37 | 42 |
| PDB IDs | 39HG | 39HH | 39HI | 39HJ |

Table S3. X-ray data collection and refinement statistics.
